## Supplementary Methods for "A novel framework for analysis of the shared genetic background of correlated traits"

### 1. Decomposing the phenotypic correlations

For any pair of traits *y*_1_ and *y*_2_, the phenotypic correlation can be written in terms of their heritabilities, *h*_1_^2^ and *h*_2_^2^, as (Falconer & Mackay 1981):

$$u_{phen}=u_{gen}\sqrt{h_{1}^{2}h_{2}^{2}}+u_{env}\sqrt{\left( 1-h_{1}^{2} \right)\left( 1-h_{2}^{2} \right)}, (M1)$$

where *u_gen_* and *u_env_* are the genetic and environmental correlations, respectively. In a similar manner, *u_gen_* can be written in terms of *w*_1_^2^ and *w*_2_^2^:

$$u_{gen}=u_{shar}\sqrt{w_{1}^{2}w_{2}^{2}}+u_{unsh}\sqrt{\left( 1-w_{1}^{2} \right)\left( 1-w_{2}^{2} \right),} (M2)$$

where *u_shar_* and *u_unsh_* are the correlations due to the SGF and UGF, respectively.

Extrapolation of Formulas (M1) and (M2) on *K* traits gives:

$$U_{phen}=\underset{genetic component}{\underbrace{\sqrt{H^{2}}U_{gen}\sqrt{H^{2}}}}+\underset{environmental component}{\underbrace{{\sqrt{I-H^{2}}U}_{env}\sqrt{I-H^{2}}}} , (M3)$$

and

$$U_{gen}=\underset{due to SGF}{\underbrace{\sqrt{W^{2}}U_{shar}\sqrt{W^{2}}}}+\underset{due to UGF}{\underbrace{\sqrt{I-W^{2}}U_{unsh}\sqrt{I-W^{2}}}}. (M4)$$

According to Model (2) of the main text and its requirements, *U_shar_* is determined in Expression (M4) as a (*K×K*) unit-rank matrix, whose elements consist only of the numbers -1 or 1 depending on the sign of the corresponding elements of *U_gen_*, *U_shar_*=*sign*(*U_gen_*); *W*^2^ is a diagonal matrix, whose *i-*th diagonal element is *w_i_*^2^; *U_unsh_* is a matrix of genetic correlations explained by the UGF; *H*^2^ is a diagonal matrix, whose *i-*th diagonal element is *h_i_*^2^; *U_env_* is an environmental correlation matrix.

Without loss of generality, Expression (M4) can be rewritten as

$$U_{gen}=W\boldsymbol{11}^{T}W+\sqrt{I-W^{2}}U_{unsh}\sqrt{I-W^{2}} , (M5)$$

where **1** is a (*K*×1) vector of units. The signs of the diagonal elements of *W* are of importance, because they indicate the directions of the SGF effects on the original traits.

The model for decomposing the correlations between the traits into components and its input/output data are presented in Figure M1.

### 2. The MaxSH method

To identify the SGF, we suggest building a new trait, SGIT, defined as a linear combination of original traits. The coefficients, *α,* of the linear combination are estimated to maximize the heritability of SGIT explained by SGF. Additionally, we build residual traits called ‘UGITs’ by regressing each original trait on the SGIT. The UGITs are the linear combinations of the original traits, too.

We developed the MaxSH method, which 1) determines the proportion of the heritability of each original trait, which is explained by the SGF; 2) calculates the *α* coefficients and 3) calculates the *γ* coefficients, to build the UGITs.

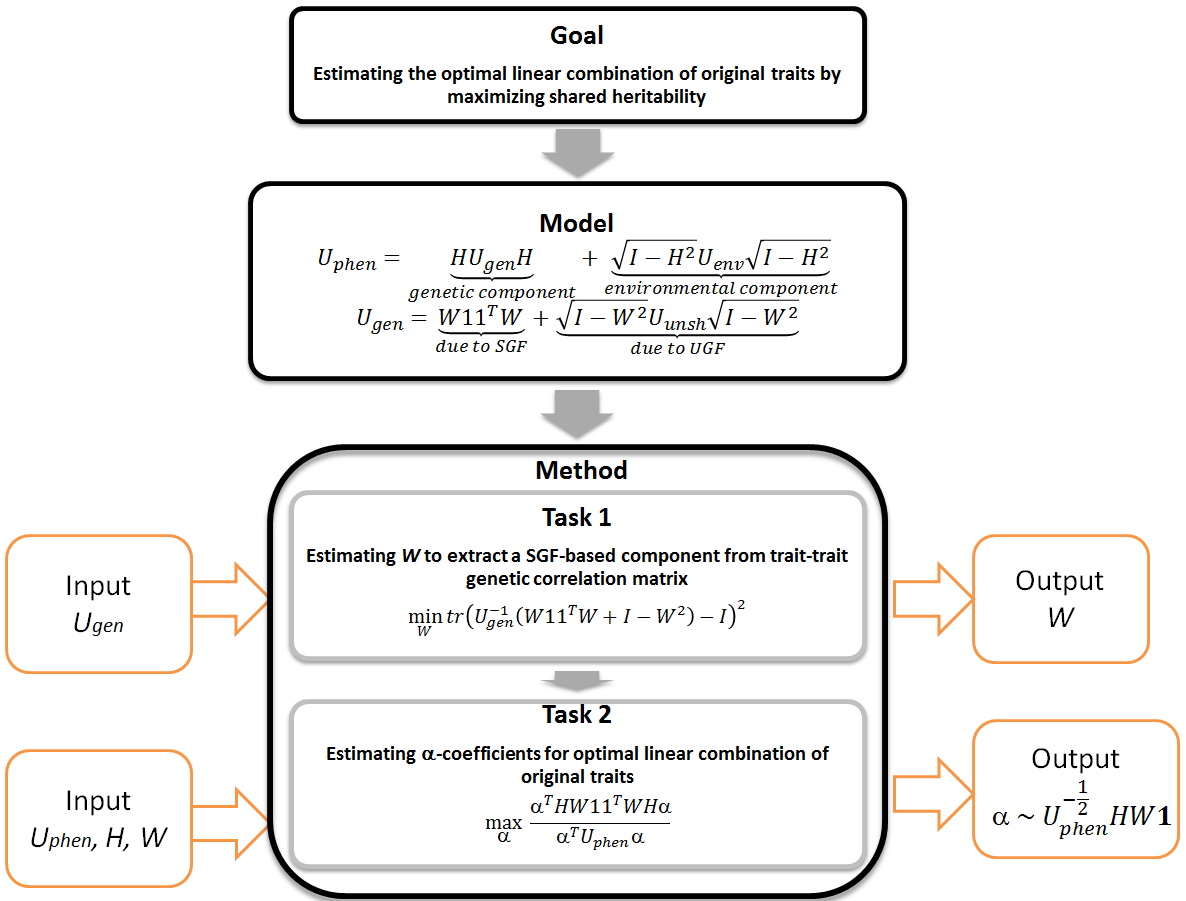

**Figure M1. A schematic of the MaxSH method for calculating the coefficients of the linear combination of the traits, SGIT.**

*Notations*. *U_phen_* and *U_gen_*: matrices of phenotypic end genotypic correlations between traits, respectively; *U_env_*: matrix of correlations caused by environmental factors; *U_unsh_*: matrix of genetic correlations caused by the UGF; *H*^2^ and *W^2^*: diagonal matrices whose diagonal elements are traits' heritabilities (*h*^2^) and their fractions explained by the SGF (*w*^2^), respectively; *α*: coefficients of SGIT, an optimal linear combination of traits; *tr*: the trace of a matrix; **1**: vector of units.

#### 2.1. The proportion of heritability explained by the SGF

First of all, we examine the *U_gen_* matrix to test whether the existence of an SGI is possible. If the genetic correlation does not significantly differ from 0 for at least one pair of traits, the existence of the SGI is impossible and our method is non-applicable. Otherwise, we transform *U_gen_* into an extremal correlation matrix by replacing the positive and negative values with -1 and 1, respectively. Next, we define a rank of the extremal matrix. If the rank is greater than 1, there is no SGF. Otherwise, an SGF exists and our method can be applied.

For estimating *W*, we use the matrix *V*, where the influence of the UGF on the covariances between the traits is minimum:

$$V=W\mathbf{11}^{T}W+I_{k}-W^{2}$$

As the optimization metric, we use a loss function, *L*_1_, which compares the matrices *U_gen_* and *V* (Konno 2010; Khodadadi & Tarami 2011):

$$L_{1}=tr\left( U_{gen}^{-1}V-I \right)^{2}.$$

#### 2.2. The *α* coefficients

After estimating *W*, we can calculate the *α* coefficients to build the SGIT defined as $\sum_{i=1}^{K} y_{i}\alpha_{i}$. According to Formulas (M3) and (M5), the phenotypic variance of the SGIT is

$${U_{phen}(SGIT)=\alpha}^{T}U_{phen}\alpha= \alpha^{T}HW\boldsymbol{11}^{T}WH\alpha+$$

$$\alpha^{T}H\sqrt{I-W^{2}}U_{unsh}\sqrt{I-W^{2}}H\alpha+ \alpha^{T}{\sqrt{I-H^{2}}U}_{env}\sqrt{I-H^{2}}\alpha,$$

and therefore the heritability of the SGIT explained by the SGF is (Oualkacha *et al.* 2012)

$$h_{SGF}^{2}(SGIT)=\frac{\alpha^{T}S\alpha}{\alpha^{T}U_{phen}\alpha}, \left( M6 \right)$$

where $S=HW\boldsymbol{11}^{T}WH$.

We analytically estimated *α* by maximizing $h_{SGF}^{2}(SGIT)$ (Exp. M6) and taking into account the fact that *α* is the first eigenvector of the matrix product (see, for example, (Mardia *et al.* 1979)):

$$\left( U_{phen}-S \right)^{-\frac{1}{2}}S\left( U_{phen}-S \right)^{-\frac{1}{2}}.$$

Then, *α* is expressed as

$$\alpha=\frac{\left( U_{phen}-HW\boldsymbol{11}^{T}WH \right)^{-\frac{1}{2}}HW1}{\sqrt{{\boldsymbol{1}^{T}WH\left( U_{phen}-HW\boldsymbol{11}^{T}WH \right)}^{-1}HW\boldsymbol{1}}}.$$

After some transformations:

$$\alpha=\frac{U_{phen}^{-\frac{1}{2}}HW\boldsymbol{1}}{\sqrt{\boldsymbol{1}^{T}WHU_{phen}^{-1}HW\boldsymbol{1}}}. (M7)$$

Note that from Expression (M7) and the properties of eigenvectors it follows that

$$\sum_{i=1}^{K} \alpha_{i}^{2}=1.$$

### 2.3. 95% CI for linear combination coefficients

The Monte Carlo approach is used to estimate the 95% confidence interval (CI) for *α*. We performed 1000 simulation repeats. In each repeat, we simulated a (*K×K*) random matrix, Ω*^noise^*, as the noise component for *U_gen_*. Each (*i,j*)-th element (*i* > *j*) of Ω*^noise^* followed a normal distribution with zero mean and standard deviation equal to the (*i,j*)-th element of the matrix of standard errors for *U_gen_*. The resulting ‘noisy’ genetic correlation matrix was obtained as *U_gen_* + Ω*^noise^*. Then, the standardized *α-*vector was calculated by building the empirical distribution of each element of the *α*-vector. 95% CI was obtained as an absolute difference between quantile values of 0.975 and 0.025 divided by 2.

### 2.4. The minimum number of traits

For effective use of the MaxSH method, it is important to find the optimum number of traits. Since the genetic correlation matrix of *K* traits is a full-rank symmetric matrix (one of the requirements of MaxSH), it can be described by *K*(*K-*1)/2 independent parameters. To avoid over-parameterization, the number of traits in analysis must not be less than *K*(*K*-1)/2. Therefore, *K* ≥ 3.

Note that *W* can be estimated analytically when *K*=3:

$$w_{i}=\frac{{u_{gen}}_{ji}{u_{gen}}_{il}}{{u_{gen}}_{jl}}.$$

The upper bound for *K* is not set. However, it is obvious that the more traits are being analyzed, the lower SGI is to be expected (small $w_{i}$). If *K* is large, we recommend to cluster the traits and estimate the correlations between the identified clusters. If these correlations are not significant, each cluster should be analyzed independently.

### 2.5. Expected genetic correlations between the SGIT and original traits

Above, we introduced $U_{gen\_cov}=HU_{gen}H$ as a (*K×K*) matrix of genetic covariances between traits. The coefficients of expected genetic correlations between the *i-*th original trait and SGIT are estimated as

$${r_{g}}_{i,\alpha}=\frac{{U_{gen\_cov}}_{i}\alpha_{i}}{\sqrt{h_{i}^{2}h_{SGIT}^{2}}},$$

where ${U_{gen\_cov}}_{i}$ is the *i*-th column of $U_{gen\_cov}$.

We can use the comparison of the expected and estimated (for example by LD score method) genetic correlations as a validity check. Moreover, we can use the expected zero genetic correlations between the UGITs and SGIT as a validity check.

### 3. The sumCOT method

To obtain GWAS summary statistics of a trait, which represents a linear combination of an arbitrary number of traits with available GWAS summary statistics, we developed the sumCOT method. It should be noted that sumCOT uses *Z*-scores obtained by any regression model. GWAS statistics could be calculated using samples with arbitrary overlaps and different sizes. When estimating the matrix of phenotypic correlations, sample overlap should be considered. Summary statistics should be quality controlled prior to performing sumCOT, matched by SNPs and effective and reference allele order.

sumCOT requires the following inputs:

1) *α*: a (*K* × 1) vector of linear combination coefficients for *k* traits.
2) *Z*: an (*M* × *K*) matrix of Z-scores calculated for *k* traits and *M* SNPs; *Z_i_* is the *i*-th column of $Z$.
3) *U_phen_*: a (*K* × *K*) matrix of phenotypic correlations estimated by any method that takes into account the sample overlap between GWASs, for example, the *Z*-score-based method proposed by Stephens (Stephens 2013).
4) *N*: an (*M* × *K*) matrix of sample sizes for *M* SNPs and *K* traits; *N_i_* is the *i*-th column of $N$.
5) *EAF*: an (*M* × 1) vector of the effective allele frequencies of *M* SNPs; *EAF* can be estimated using a reference sample or obtained from any of the GWASs used in analysis.

Let us denote the resulting linear combination as LCT (linear combination trait).

#### Step by step

1) Estimate the expected phenotypic variance of the LCT: $Var(LCT)=\sum\left[ (\alpha\bigotimes\alpha)\circ U_{phen} \right]$, where $\bigotimes$ is an outer product.
2) For each GWAS, standardize SNP effect sizes and their standard errors using *Z* scores and sample sizes (note that these are not yet scaled on SNP variance: ${SE}_{i}^{s,u}=\sqrt{1/{({Z_{i}}^{2}+N_{i})}}$ and $\beta_{i}^{s,u}=Z_{i}*{SE}_{i}^{s,u}$ (*i* = 1, …, *K*).
3) For each SNP, combine unscaled SNP effects into raw (unstandardized, unscaled) LCT effect sizes, using *α* as weights: $\beta_{LCT,SNP}^{u,u}=\beta_{SNP}^{s,u}\times\alpha$, where $\times$ is an inner product and $\beta_{SNP}^{s,u}$ is a vector of standardized effect sizes for the given SNP and *k* traits. As a results, we have $\beta_{LCT}^{u,u}$, an (*M* × 1) vector of standardized effects sizes for *M* SNPs.
4) For each SNP, estimate the standard error of the unstandardized, unscaled LCT effect size. ${SE}_{LCT,SNP}^{u,u}=\sqrt{\sum[(\alpha\otimes\alpha)\circ U_{phen}\boldsymbol{\circ}\left( {SE}_{SNP}^{s,u}\otimes{SE}_{SNP}^{s,u}) \right]}$, where $\otimes$ is an outer product and ${SE}_{SNP}^{s,u}$ is a vector of standardized standard errors for the given SNP and *K* traits. As a results, we have ${SE}_{LCT}^{u,u}$, an (*M* × 1) vector of standardized standard errors for *M* SNPs. Note the $U_{phen}$ term inflates the standard errors to account for the correlations between GWAS statistics (i.e. sample overlap).
5) Standardize the LCT effect sizes and standard errors: $\beta_{LCT}^{s,u}={\beta_{LCT}^{u,u}}/\sqrt{Var(LCT)}$ and ${SE}_{LCT}^{s,u}={{SE}_{LCT}^{u,u}}/\sqrt{Var(LCT)}$.
6) Estimate the effective sample size of the LCT*:* $N_{LCT}=median(1/{({{SE}_{LCT}^{s,u}}^{2})})$*.* This step is derived from Equation (1) from (Winkler *et al.* 2014), which estimates the expected sample size of the trait from standard errors: $\sqrt{N}=median(1/\sqrt{Var(SNP}))*\sqrt{Var\left( Y \right)}*median\left( 1/{SE} \right)$. In our case, the expected phenotypic variance of the LCT is equal to one, and the variance of SNPs is equal to one, since the unscaled standard errors are used ($var\left( Y \right)=1$ and $var\left( SNP \right)=1$).
7) Transform unscaled LCT effect sizes and standard errors on SNP variance: $\beta_{LCT}^{s,s}={\beta_{LCT}^{s,u}}/\sqrt{Var(SNP)}$ and ${SE}_{LCT}^{s,s}={{SE}_{LCT}^{s,u}}/\sqrt{Var(SNP)}$, where $Var\left( SNP \right)= 2*EAF*\left( 1-EAF \right)$.
8) Estimate the corresponding P values using the Wald test ($Z_{LCT}={\beta_{LCT}^{s,s}}/{{SE}_{LCT}^{s,s}}$).

### 4. Genetic and phenotypic correlation matrices for simulation studies

We generated random matrices, *U_unsh_* and *U_env_*, for each simulation iteration using the following algorithm:

1. Generate *K*(*K*-1)/2 elements of the upper triangular matrix using a generator of uniformly distributed random numbers in the interval (- *d*_1_, *d*_1_) for *U_unsh_* and (- *d*_2_, *d*_2_) for *U_env_*.
2. Randomly replace *s***K*(*K*-1) elements of the upper triangular matrix in *U_unsh_* by zeros (for *U_env_* , this procedure is not applied).
3. Construct the square matrices *U_unsh_* and *U_env_* from the upper triangular matrix form.

Having generated the matrix *U_unsh_* and using the decomposition model (see Fig. M1, box ‘Model’), we can calculate *U_gen_.* In each simulation iteration, *U_unsh_* and *U_gen_* should be checked against the following criteria:

(1) Both matrices should be positively defined.

(2) Each element of *U_gen_* had to be more than or equal to an assigned threshold (0.1).

(3) *U_gen_* must have an SGI, while *U_unsh_* must have none.

If at least one of the criteria was not met, we did not take into account this iteration. If all criteria were met, we generated $U_{phen}$ using the model (see Fig. M1, box ‘Model’). The matrices *U_gen_* and *U_phen_*, and heritabilities were used as input data in the GIP and MaxSH approaches.
