## Supplementary Results for "A novel framework for analysis of the shared genetic background of correlated traits"

### Simulation results

Under different scenarios, we designed simulations to assess the performance of MaxSH. We (1) assessed the accuracy of *w* estimates with respect to the loss function, (2) assessed the proportion of shared heritability in the total heritability of the SGIT (the *Q*-value) with respect to the loss function, and (3) compared the analytically predicted total/shared heritabilities of two traits: SGIT and the first genetically independent phenotype GIP1.

All results are presented in Supplementary Figures S1-18. Each figure consists of four panels (a-d). Panel (a) is the plot of ${\Delta W=\left( \frac{\text{W}\text{0}\text{-West}}{\text{W}\text{0}} \right)}^{2}$versus the loss function. Panel (b) is the plot of the total heritability of the SGIT versus the total heritability of GIP1. Panel (c) is the plot of the *Q*-value versus the loss function. Panel (d) is the plot of the shared heritability of the SGIT versus the shared heritability of GIP1. Different colors represent different *W^2^* values. The values of other parameters are presented in the main title of the figure. “All h2 are different” means that h^2^ varied from 0.3 to 0.8.

**Figures S1-6. Results of simulations for three traits (K=3).** All details are in the text.

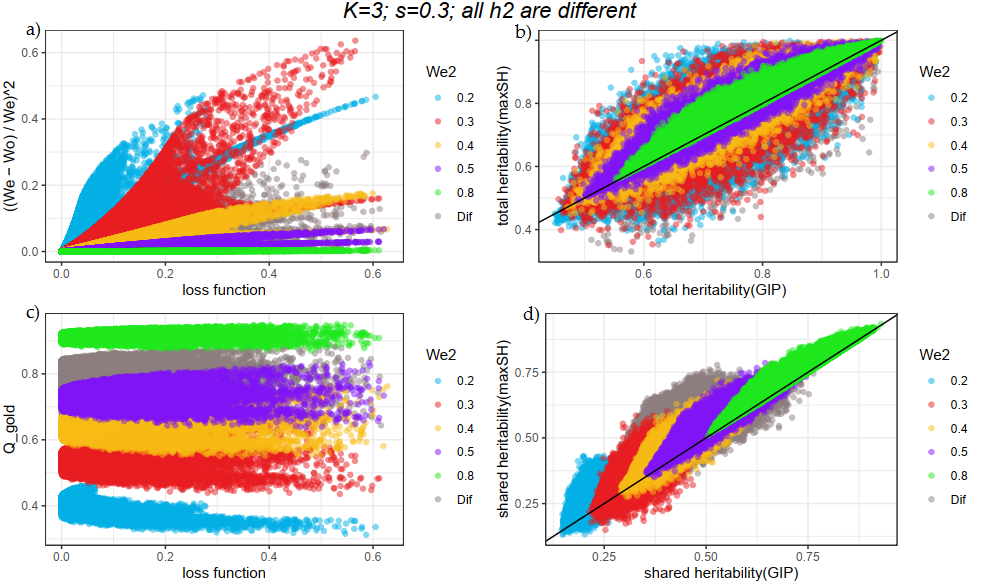

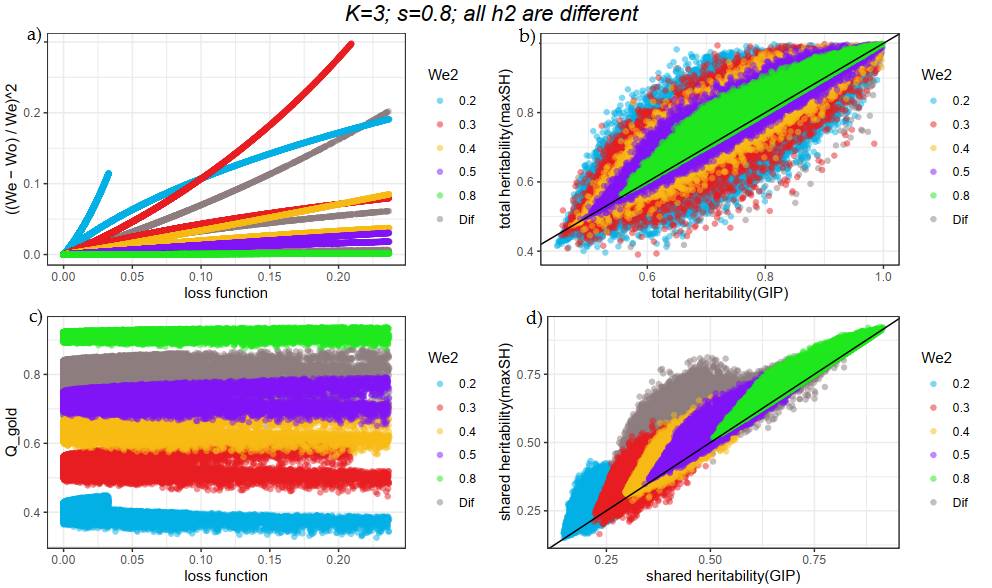

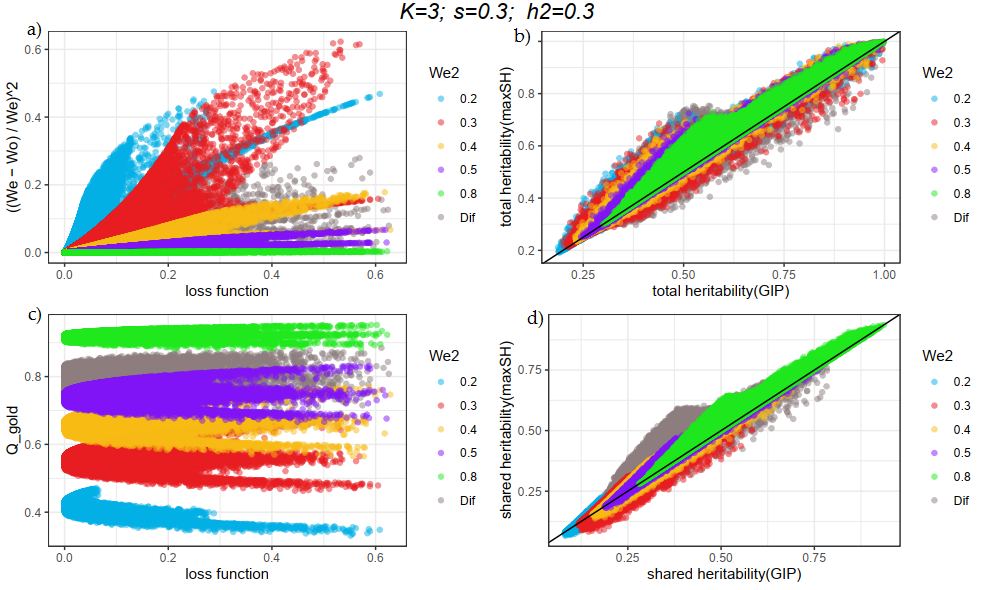

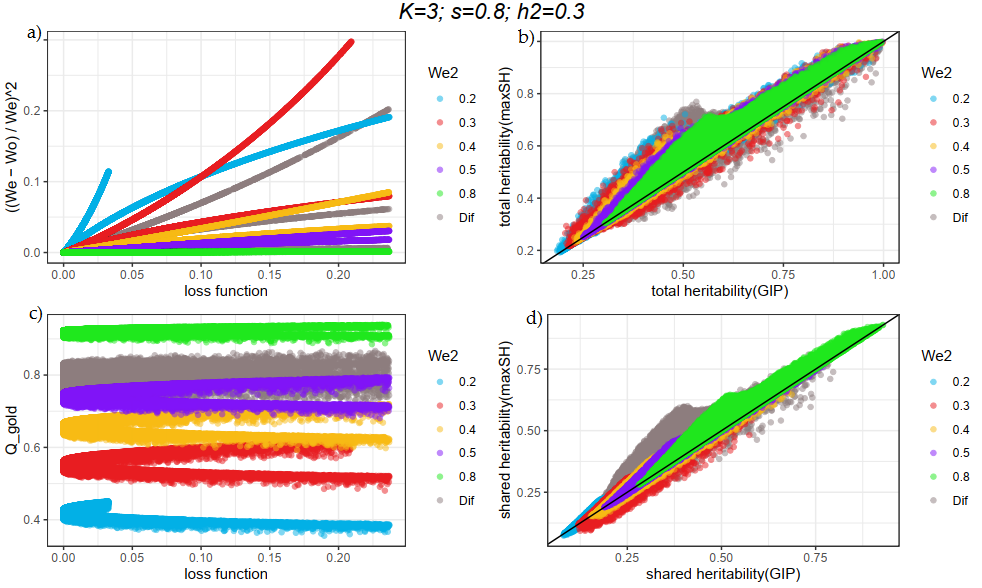

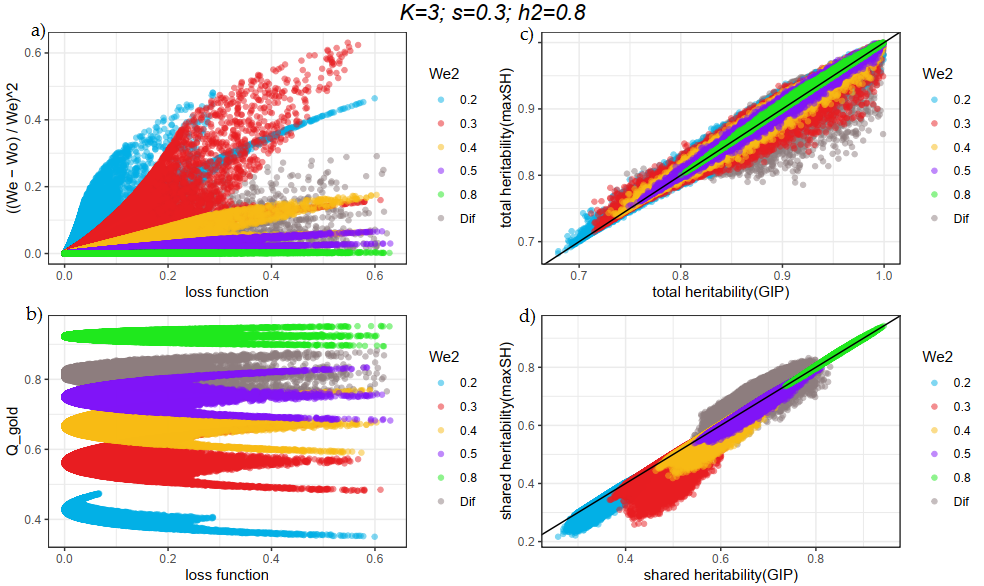

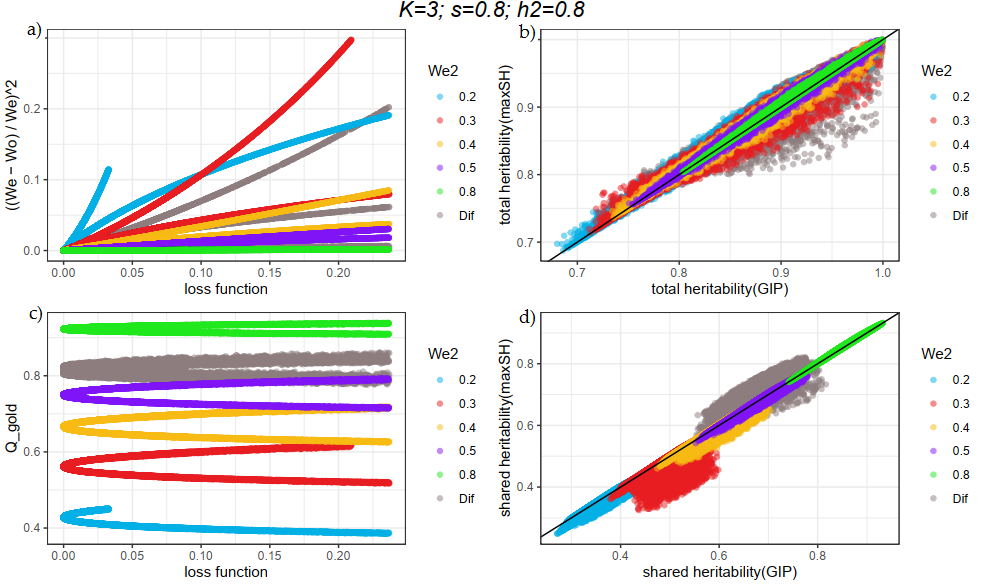

**Figures S7-12. Results of simulations for four traits (K=4).** All details are in the text.

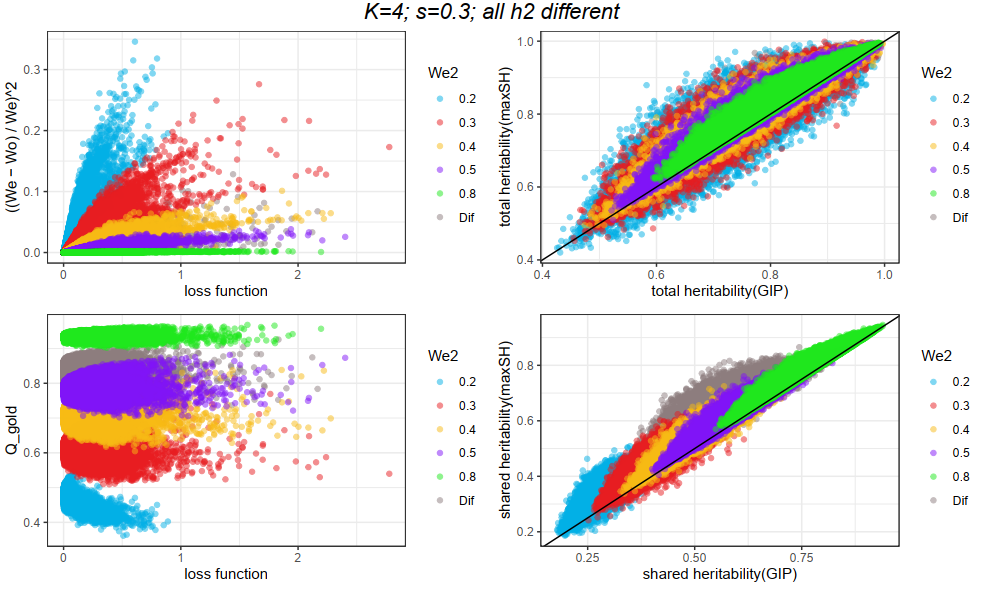

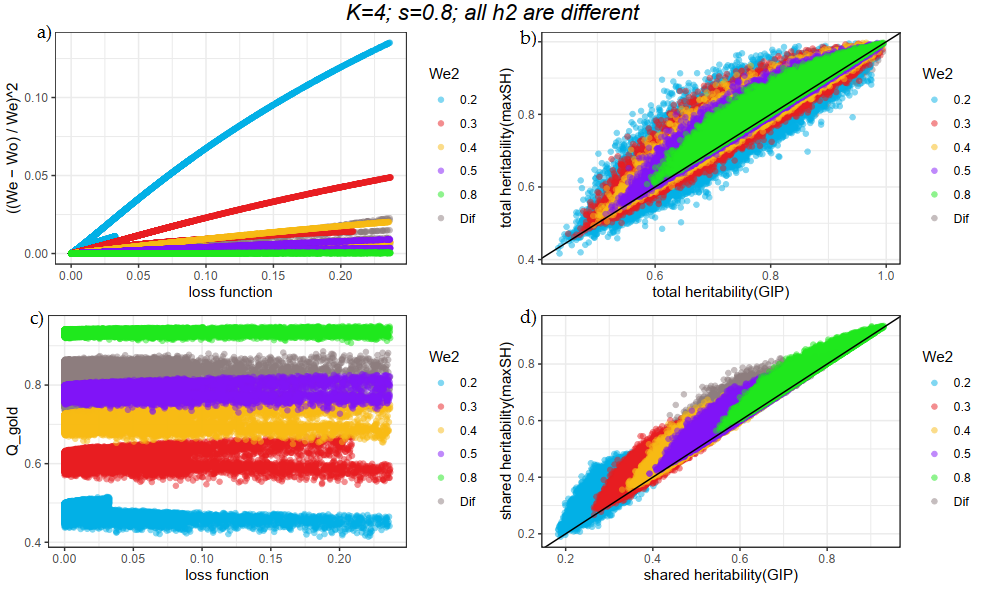

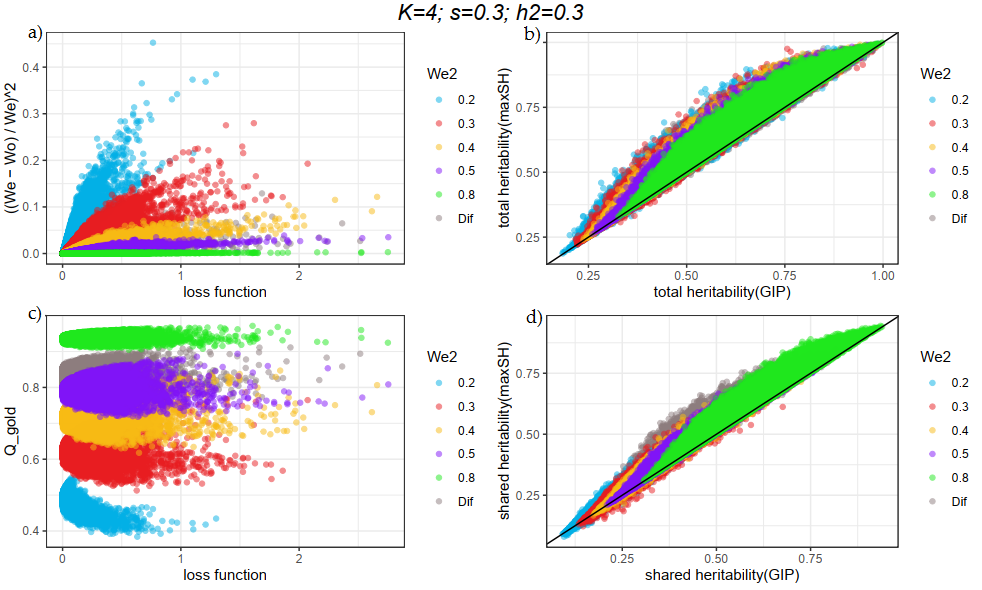

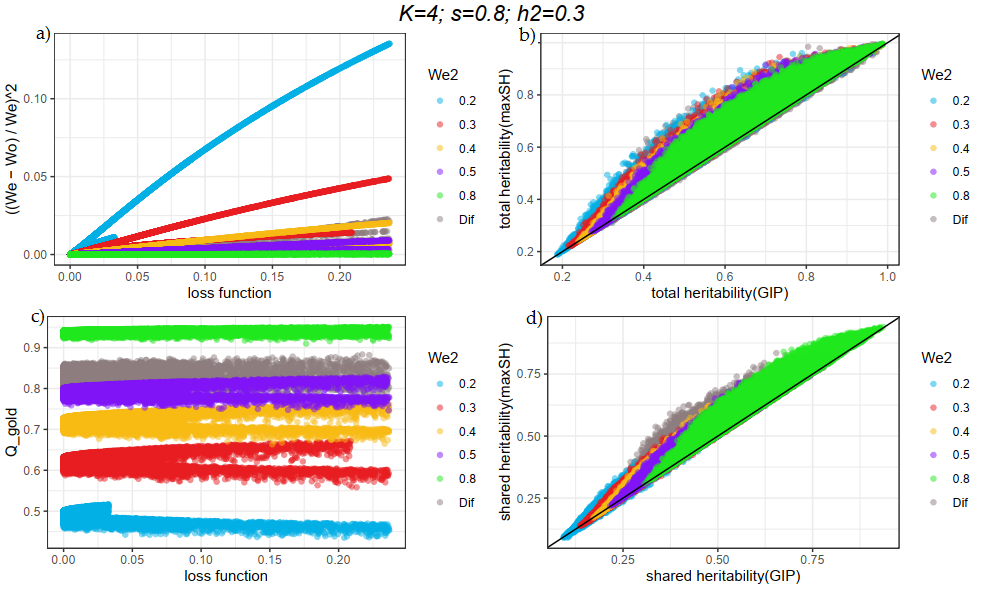

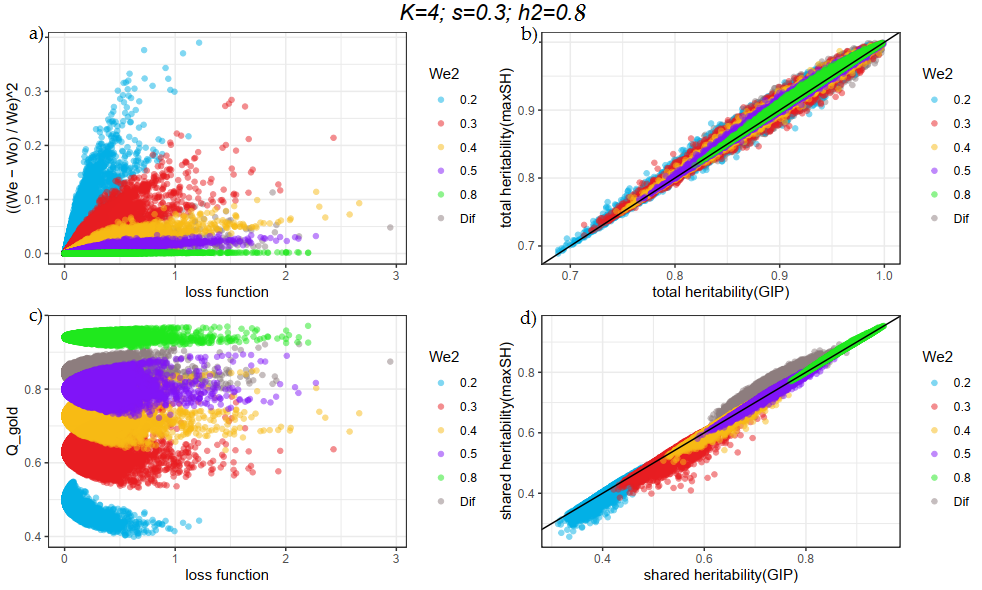

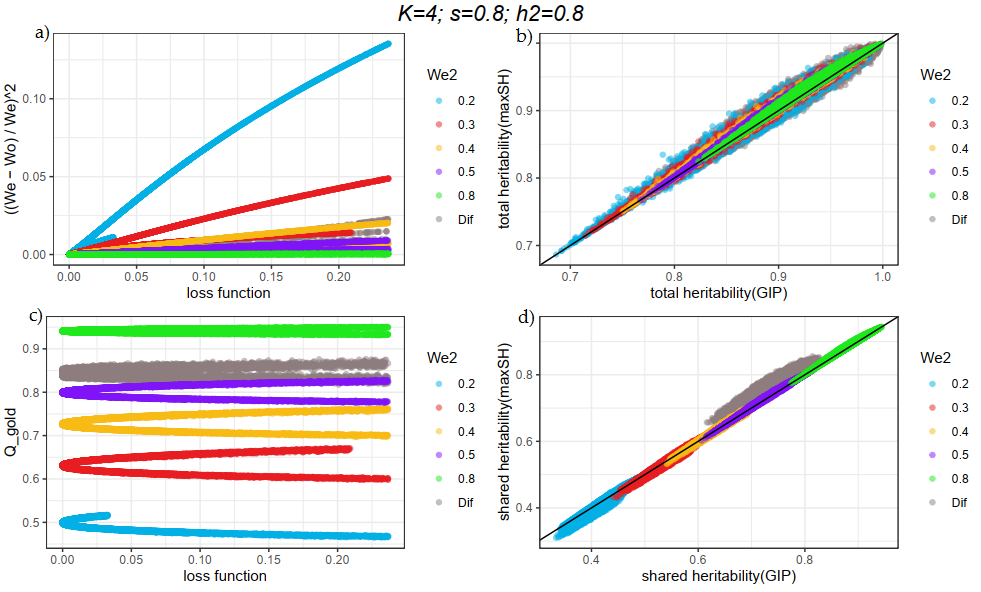

**Figures S13-18. Results of simulations for five traits (K=5).** All details are in the text**.**

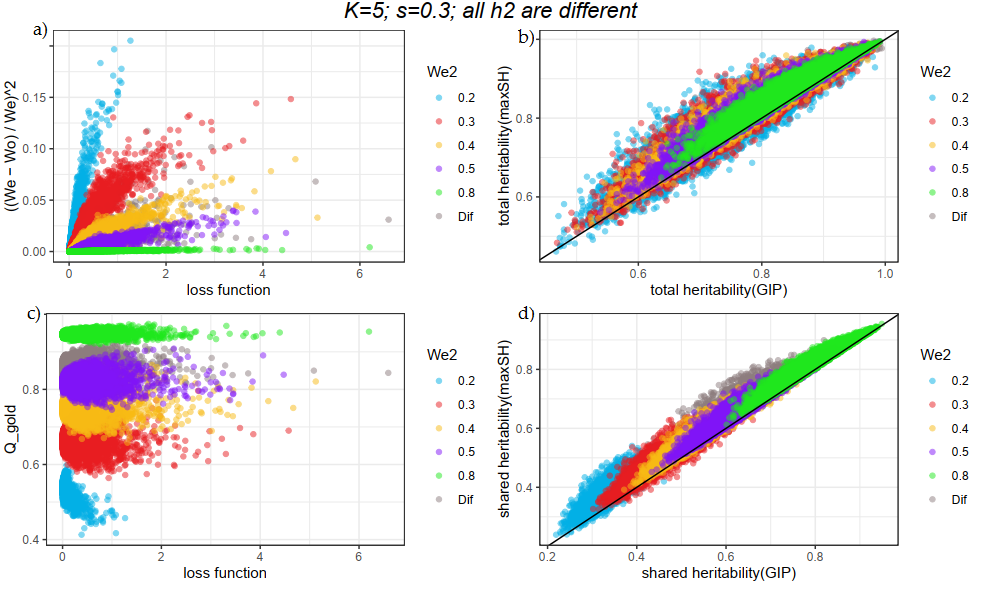

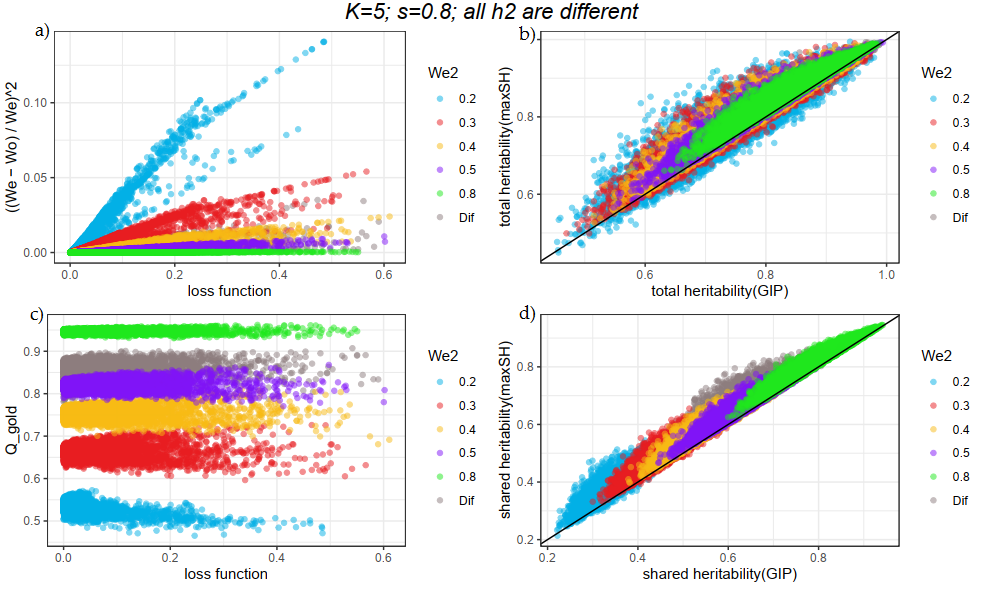

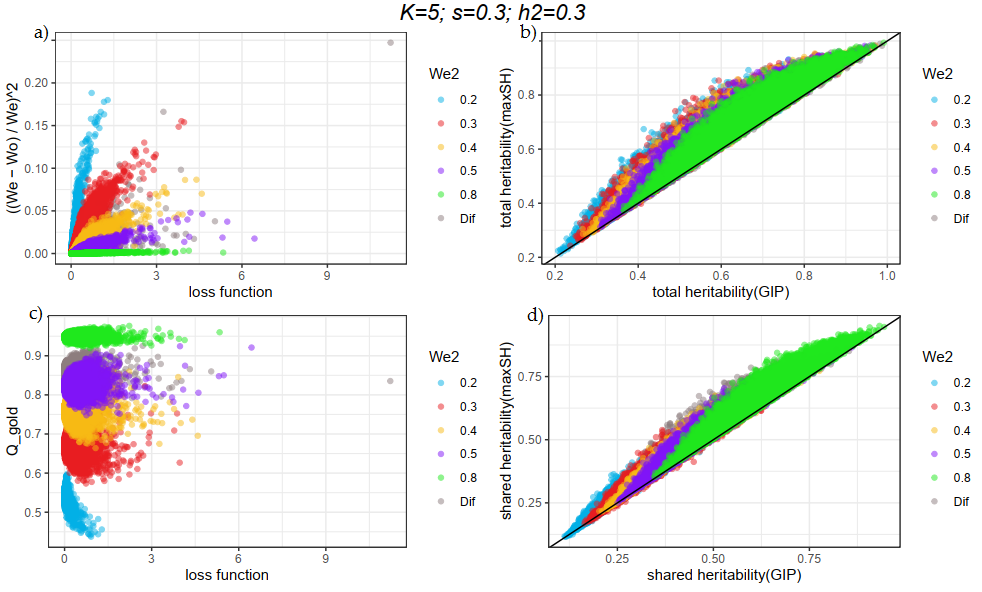

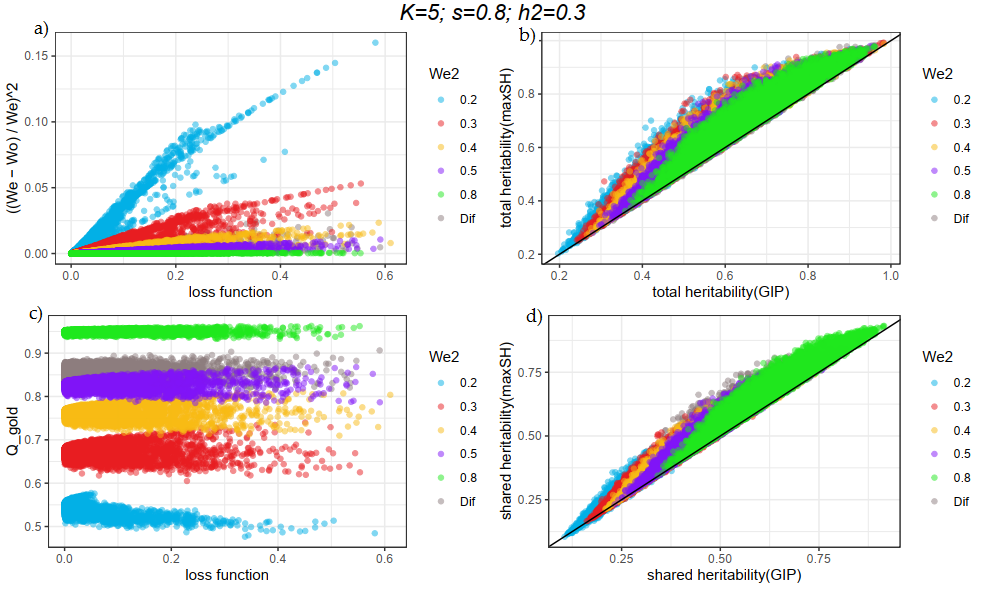

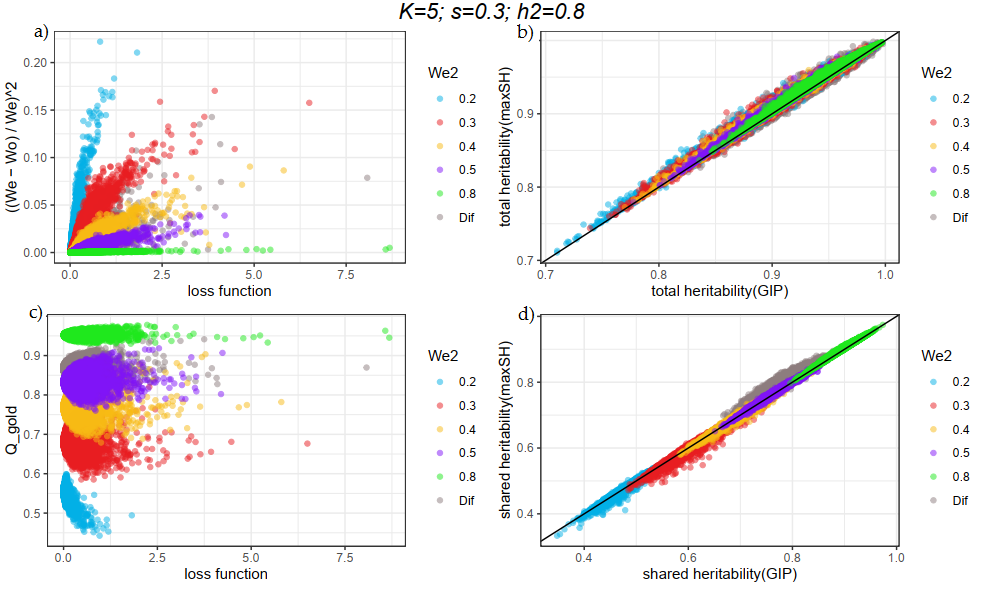

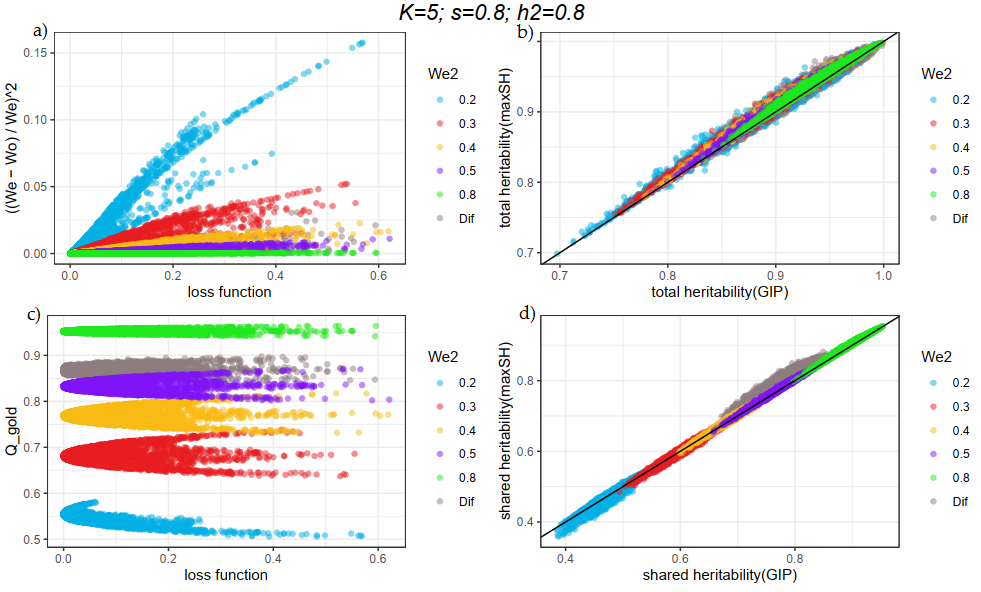

### Real data assessment

We applied the SHAHER framework to three datasets: anthropometric (five traits), psychometric (four traits) and lipids (three traits). The results for anthropometric traits are presented in the main text. We confirmed that SGI exists for each dataset. For each dataset, there was no SGI between UGITs, which was up to expectation. The full results of clumping for original traits, SGIT and UGITs are presented in Supplementary Tables 3a-c. The number of significant loci for each dataset is presented in Table S1. The full results of the DEPICT analyses are presented in Supplementary Tables 4-6. The number of significantly enriched sets and tissues (FDR<0.05) is presented in Table S2.

**Table S1.** Number of significant loci for each trait at p-values < 5×10^-8^

| Trait name | Number of significant loci |
| --- | --- |
| Anthropometric traits |  |
| BMI | 344 |
| Weight | 394 |
| Hip | 315 |
| Waist | 239 |
| Fat | 313 |
| SGIT | 337 |
| BMI UGIT | 262 |
| Weight UGIT | 212 |
| Hip UGIT | 100 |
| Waist UGIT | 79 |
| Fat UGIT | 52 |
| PGC |  |
| BIP | 14 |
| MDD | 3 |
| SCZ | 96 |
| Happiness | 0 |
| SGIT | 67 |
| BIP UGIT | 3 |
| MDD UGIT | 3 |
| SCZ UGIT | 2 |
| Happiness UGIT | 1 |
| Lipid concentrations |  |
| LDL | 87 |
| Triglycerides | 78 |
| Cholesterol | 103 |
| SGIT | 99 |
| LDL UGIT | 48 |
| Triglycerides UGIT | 68 |
| Cholesterol UGIT | 59 |

**Table S2.** Number of significantly enriched sets and tissues for DEPICT analysis (FDR<0.05)

|  | Number of  enriched sets | Number of  enriched tissues |
| --- | --- | --- |
| Anthropometric traits |  |  |
| BMI | 192 | 22 |
| Weight | 814 | 0 |
| Hip | 825 | 0 |
| Waist | 4 | 19 |
| Fat | 68 | 19 |
| SGIT | 246 | 21 |
| BMI UGIT | 1608 | 47 |
| Weight UGIT | 0 | 45 |
| Hip UGIT | 58 | 9 |
| Waist UGIT | 40 | 1 |
| Fat UGIT | 2 | 4 |
| PGC |  |  |
| BIP | 0 | 0 |
| SCZ | 0 | 0 |
| SGIT | 9 | 23 |
| Lipid concentrations |  |  |
| LDL | 354 | 3 |
| Triglycerides | 394 | 11 |
| Cholesterol | 359 | 3 |
| SGIT | 545 | 3 |
| LDL UGIT | 227 | 28 |
| Triglycerides UGIT | 531 | 37 |
| Cholesterol UGIT | 286 | 0 |

#### Psychometric traits

Figure S19-A demonstrates the genetic correlations for original psychometric traits and their UGITs. All the original traits were positively correlated with *r* > 0.21. The minimal correlation of SGIT with the original traits was observed for happiness (*r* = 0.36), and the maximal correlation was 0.94 with BIP. For UGITs, we did not observe significant genetic correlations with SGIT, nor did we find additional SGI. The heritabilities of UGITs varied from 0.06 to 0.11.

Boxplots of the dependence between the SGIT p-value (Figure S19-B) and the number of original traits significantly associated with the locus show that the higher the number, the lower SGIT p-values.

The joint clumping of 9 traits (4 original traits, SGIT and 4 UGITs) revealed 135 significantly (p-value < 5×10^-8^) associated loci (see Supplementary Table 3b). The SGIT was associated with 67 loci, of which 24 were not significantly associated with the original traits and were thus considered new. The number of loci associated with the original traits varied strongly (from 0 for Happiness to 96 for SZC), and so the heatmap of relative overlapping between loci (Figure S19-C) is not informative. The joint clumping of only original traits revealed 107 loci, out of which 62 could not be detected by analysis of the SGIT or UGITs. The joint clumping of the SGIT and UGITs revealed 73 loci, of which 28 could not be detected by analysis of the original traits and were thus considered new.

The results of gene set and tissue enrichment analyses are presented only for the BIP and SCZ traits, and SGIT (an analysis is applicable if the number of significant loci is greater than 10) in Supplementary Table 5. In gene set and tissue enrichment analyses, significant results were observed only for the SGIT, although the number of hits was not the highest among the traits (SCZ has 96 significant loci). We observed 23 significantly enriched tissues related to the nervous system and 9 gene sets associated with the nervous system development. Thus, we concluded that the SGIT was genetically more homogenous than the original traits.

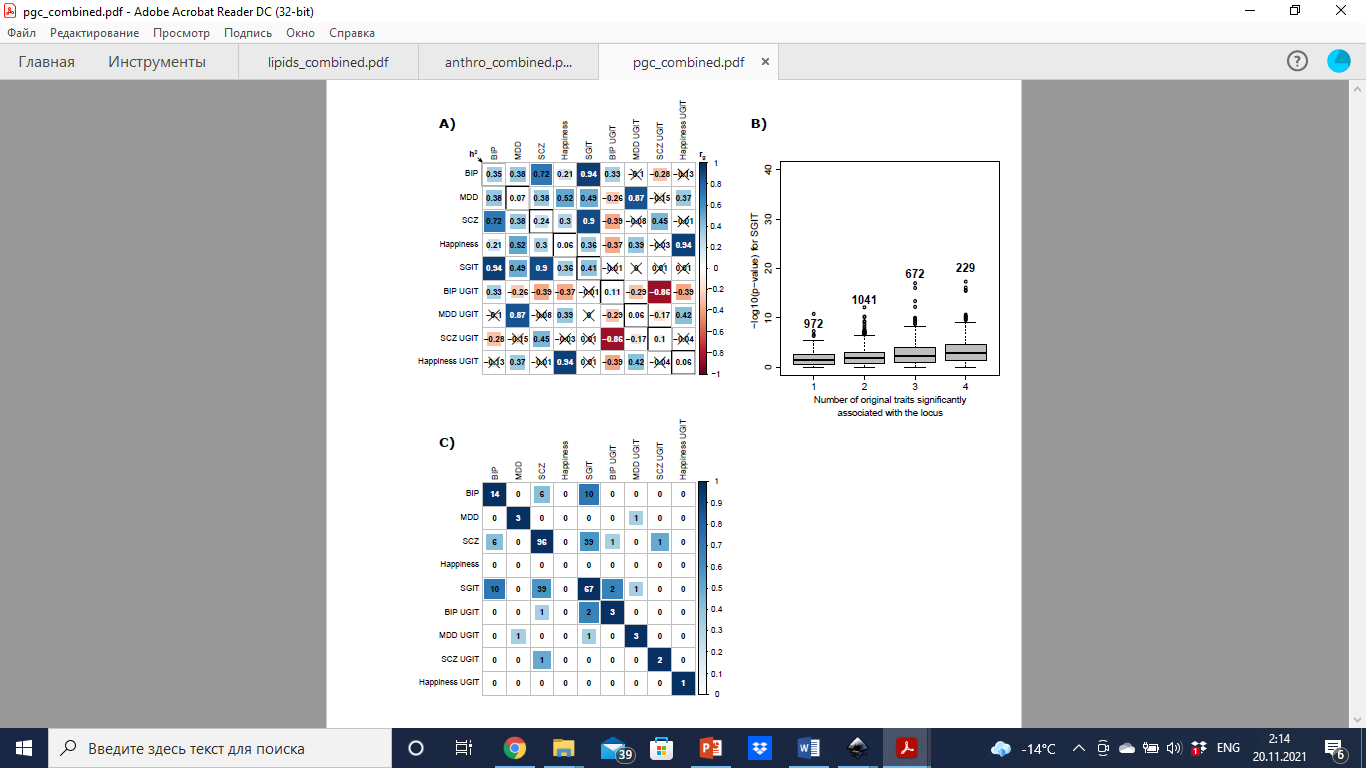

**Figure S19.** **The results of the application of SHAHER on four psychometric traits.** A) The heatmap of genetic correlations between the original, SGI and UGI traits. The number, color strength and size of the squares in the matrix show the values of the correlation coefficients between the traits. The diagonal elements represent heritabilities. Crossed out values indicate insignificant correlations. B) Boxplots of –log_10_(p-value) for the SGIT with respect to the number of the original traits significantly associated with the locus. The number at the top of the boxplot corresponds to the number of significant SNPs. C) The heatmap of the numbers of overlapping loci between traits. The numbers in the cells represent the absolute numbers of overlapping loci. The color strength and size of the squares in the cells show the relative scaled number of overlapping loci (on the scale from 0 to 1). The diagonal elements represent the number of loci found for every trait.

#### Lipid traits

We applied SHAHER to three lipid traits. It should be noted that this is the minimum number of traits that could be used in our framework. Figure S20-A demonstrates the genetic correlations between all pairs of the original lipid traits and their UGITs. All the original traits were correlated positively with r>0.42. The genetic correlations between SGIT and cholesterol / LDL levels were almost equal to one. The genetic correlation between triglycerides and SGIT was lower (0.74), but much higher than the correlations between triglycerides and cholesterol / LDL levels (0.44 and 0.42, respectively). We observed no significant genetic correlation between SGIT and UGITs and there were no additional SGIs between them either. The heritabilities of the UGITs varied from 0.10 to 0.17.

The boxplots of the dependence of the SGIT p-value (Figure S20-B) on the number of the original traits significantly associated with the locus show that the higher this number, the lower the p-value of the SGIT.

The SGIT was genome-wide significantly associated (p-value < 5×10^-8^) with 103 loci (see Supplementary Table 3c), of which two were not significantly associated with the original traits and were thus considered new.

The heatmap represented on Figure S20-C corresponds to the results of joint clumping and shows the number of overlapping between traits from the dataset.

The results of gene set and tissue enrichment analyses are presented in Supplementary Tables 6. The overlap between the gene sets is depicted in Figure S20-D. The SGIT had the largest number of significantly enriched gene sets among all the traits (N=545). Although genetic correlations between UGITs and SGIT did not reach significance, overlapping between the gene sets was quite high.

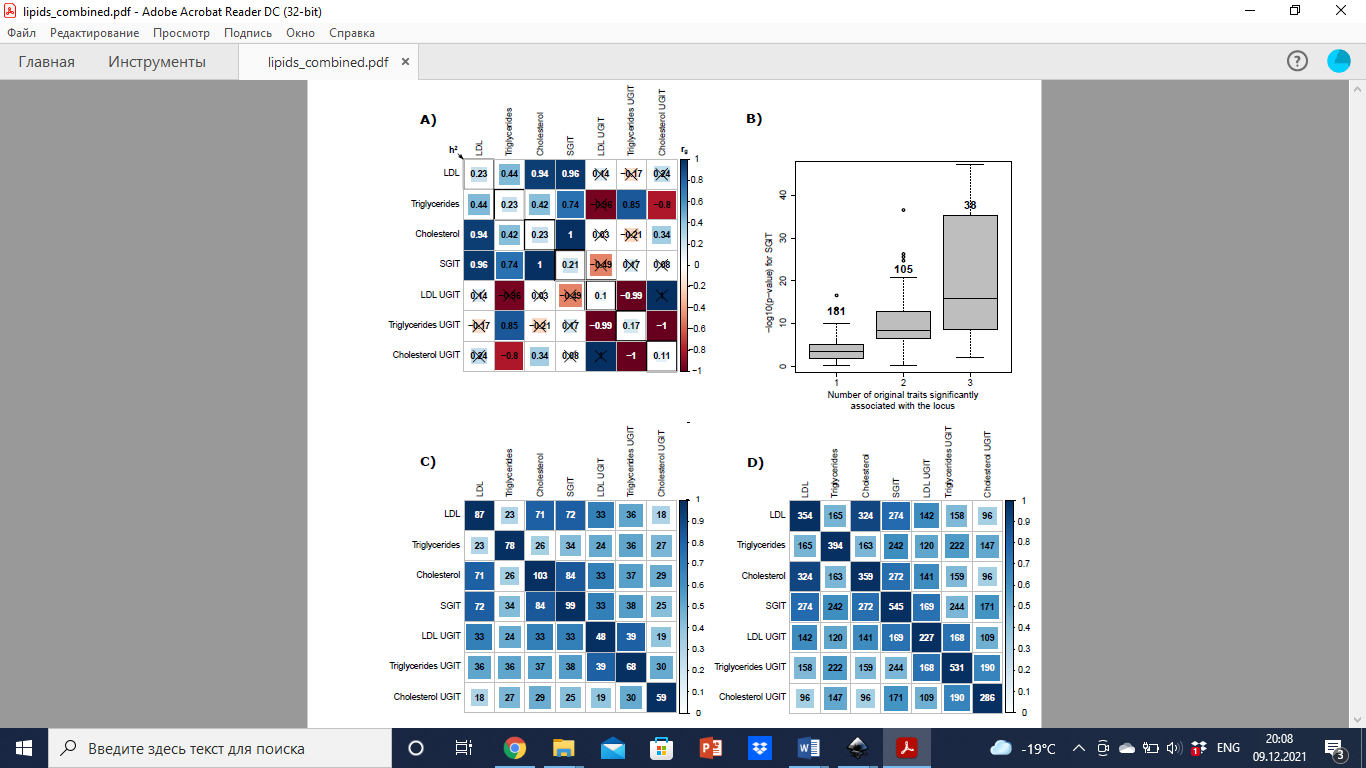

**Figure S20.** **The results of the application of SHAHER on three lipid traits.** A) The heatmap of genetic correlations between the original, SGI and UGI traits. The number, color strength and size of the squares in the matrix show the values of the correlation coefficients between the traits. The diagonal elements represent heritabilities. Crossed out values indicate insignificant correlations. B) Boxplots of –log_10_(p-value) for the SGIT with respect to the number of the original traits significantly associated with the locus. The number at the top of the boxplot corresponds to the number of significant SNPs. C) The heatmap of the numbers of overlapping loci between traits. The numbers in the cells represent the absolute numbers of overlapping loci. The color strength and size of the squares in the cells show the relative scaled number of overlapping loci (on the scale from 0 to 1). The diagonal elements represent the number of loci found for every trait. D) The heatmap of the numbers of overlapping gene sets between traits. The color strength and size of the squares in the cells show the relative scaled number of overlapping gene sets (on the scale from 0 to 1). The diagonal elements represent the number of gene sets found for every trait.
